# Membrane tethering potency of Rab-family small GTPases is defined by the C-terminal hypervariable regions

**DOI:** 10.1101/2020.06.28.176792

**Authors:** Sanae Ueda, Naoki Tamura, Joji Mima

## Abstract

Membrane tethering is a crucial step to determine the spatiotemporal specificity of secretory and endocytic trafficking pathways in all eukaryotic endomembrane systems. Recent biochemical studies by a chemically-defined reconstitution approach reveal that, in addition to the structurally-diverse classic tethering factors such as coiled-coil tethering proteins and multisubunit tethering complexes, Rab-family small GTPases also retain the inherent membrane tethering functions to directly and physically bridge two distinct lipid bilayers by themselves. Although Rab-mediated membrane tethering reactions are fairly efficient and specific in the physiological context, its mechanistic basis is yet to be understood. Here, to explore whether and how the intrinsic tethering potency of Rab GTPases is controlled by their C-terminal hypervariable region (HVR) domains that link the conserved small GTPase domains (G-domains) to membrane anchors at the C-terminus, we quantitatively compared tethering activities of two representative Rab isoforms in humans (Rab5a, Rab4a) and their HVR-deleted mutant forms. Strikingly, deletion of the HVR linker domains enabled both Rab5a and Rab4a isoforms to enhance their intrinsic tethering potency, exhibiting 5-to 50-fold higher initial velocities of tethering for the HVR-deleted mutants than those for the full-length, wild-type Rabs. Furthermore, we revealed that the tethering activity of full-length Rab5a was significantly reduced by the omission of anionic lipids and cholesterol from membrane lipids and, however, membrane tethering driven by HVR-deleted Rab5a mutant was completely insensitive to the headgroup composition of lipids. Reconstituted membrane tethering assays with the C-terminally-truncated mutants of Rab4a further uncovered that the N-terminal residues in the HVR linker, located adjacent to the G-domain, are critical for regulating the intrinsic tethering activity. In conclusion, our current findings establish that the non-conserved, flexible C-terminal HVR linker domains define membrane tethering potency of Rab-family small GTPases through controlling the close attachment of the globular G-domains to membrane surfaces, which confers the active tethering-competent state of the G-domains on lipid bilayers.

## Introduction

All eukaryotic cells, from a unicellular yeast to human cells, organize the complex but highly-regulated endomembrane systems, in which diverse cellular components including proteins and lipids are selectively delivered to their correct destinations, such as subcellular organelles, the plasma membrane, or the extracellular space, through secretory and endocytic trafficking pathways (Bonifacino and Glick, 2004). Membrane tethering is a reversible process of the initial physical contact between membrane-bound, cargo-loaded transport carriers (e.g., secretory and endocytic vesicles) and their target subcellular compartments (Pfeffer, 1999; Waters and Pfeffer, 1999; Waters and Hughson, 2000). The process of membrane tethering is vital for determining the spatiotemporal specificity of intracellular membrane trafficking, before the irreversible final steps of membrane docking and fusion mediated by SNARE-family proteins (Jahn and Scheller, 2006), which are another critical layers to confer the fidelity of membrane trafficking (McNew et al., 2000; Parlati et al., 2002; Izawa et al., 2012; Furukawa and Mima, 2014). A large body of prior studies on membrane tethering or vesicle tethering (or capture) have identified a number of the protein components essential for membrane tethering (Yu and Hughson, 2010; Kuhlee et al., 2015; Cheung and Pfeffer, 2016; Spang, 2016; Witkos and Lowe, 2016; Gillingham and Munro, 2019), which include the Uso1/p115 coiled-coil protein (Sapperstein et al., 1995; Sapperstein et al., 1996; Barlowe, 1997; Cao et al., 1998), golgin-family coiled-coil proteins (Drin et al., 2008; Wong and Munro, 2014; Cheung et al., 2015), the EEA1 coiled-coil protein (Murray et al., 2016), and a diversified set of multisubunit tethering complexes, such as the HOPS complex (Price et al., 2000; Stroupe et al., 2009; Hickey and Wickner, 2010; Ho and Stroupe, 2015; Ho and Stroupe, 2016), the exocyst complex (TerBush et al., 1996; Guo et al., 1999; Rossi et al., 2020), the COG complex (Ungar et al., 2002; Zolov and Lupashin, 2005; Shestakova et al., 2006), the Dsl1 complex (Reilly et al., 2001; Zink et al., 2009; Ren et al., 2009), the GARP complex (Conibear and Stevens, 2000; Perez-Victoria et al., 2008; Perez-Victoria and Bonifacino, 2009), and the TRAPP complex (Sacher et al., 2001; Cai et al., 2005; Cai et al., 2007). It is noteworthy that, in addition to these miscellaneous, sequentially- and structurally-diverse classic tethering factors, our recent reconstitution studies have established the inherent tethering functions of human Rab-family small GTPases (Tamura and Mima, 2014; Inoshita and Mima, 2017; Mima, 2018; Segawa et al., 2019), following the pioneering work of Merz and colleagues, which reported for the first time the intrinsic tethering activity of endosomal Ypt/Rab-family proteins in the yeast *Saccharomyces cerevisiae* (Lo et al., 2012).

Using the chemically-defined reconstitution system with purified proteins of putative membrane tethers or tethering factors and synthetic liposomes for a model lipid membrane, which is known as the most valid experimental approach to investigating whether or not the protein components of interest act as a *bona fide* membrane tether (Brunet and Sacher, 2014; Mima, 2018), comprehensive analyses of human Rab-family GTPases demonstrated their intrinsic membrane tethering potency to physically link two distinct lipid bilayers by themselves, even in the absence of any other tethering factors previously identified (Tamura and Mima, 2014; Inoshita and Mima, 2017; Mima, 2018; Segawa et al., 2019). Experimental evidence from the reconstitution studies further confirmed the efficiency and specificity of Rab-mediated membrane tethering in the physiological context: (1) A number of representative human Rab-family isoforms can efficiently drive tethering at a physiologically-relevant level of the Rab protein densities on membrane surfaces (Tamura and Mima, 2014; Inoshita and Mima, 2017; Segawa et al., 2019); (2) reversible membrane tethering is exclusively mediated by *trans*-assembly of the membrane-anchored forms of Rab proteins (Tamura and Mima, 2014; Inoshita and Mima, 2017; Segawa et al., 2019); (3) efficient tethering can be driven by specific heterotypic combinations of different Rab isoforms, such as the pair of Rab1a and Rab9a (Segawa et al., 2019); and (4) Rab11a and its cognate effector proteins, class V myosins, specifically cooperate to trigger rapid membrane tethering in a GTP-dependent manner (Inoshita and Mima, 2017). However, in spite of these research advances, the mechanistic basis of Rab-driven membrane tethering reactions remains poorly understood. In this study, by quantitatively analyzing the membrane tethering capacities of human endosomal Rabs (Rab5a and Rab4a) and their mutant forms lacking the C-terminal hypervariable region (HVR) domains that link a conserved small GTPase domain to a membrane anchor at the C-terminus, we uncovered that deletion of the HVR linkers allows Rab proteins to enhance their intrinsic tethering potency, establishing the essential role of the non-conserved flexible HVR linkers in controlling Rab-mediated membrane tethering.

## Materials and Methods

### Protein expression and purification

Bacterial expression vectors for the full-length proteins of human Rab5a (amino acid residues, Met1-Asn215; UniProtKB: P20339) and Rab4a (amino acid residues, Met1-Cys218; UniProtKB: P20338) and their mutant forms lacking the HVR linkers, Rab5aΔHVR (amino acid residues, Met1-Pro182) and Rab4aΔHVR (amino acid residues, Met1-Leu175), were constructed using a pET-41 Ek/LIC vector kit (Novagen) (**Figure 1**), as described (Tamura and Mima, 2014; Inoshita and Mima, 2017; Segawa et al., 2019). DNA fragments encoding these wild-type and mutant proteins of human Rabs and the additional sequences for a human rhinovirus 3C protease-cleavage site (Leu-Glu-Val-Leu-Phe-Gln-Gly-Pro) at the N-terminus and for a polyhistidine tag (His12) at the C-terminus were amplified by PCR using KOD-Plus-Neo polymerase (Toyobo) and Human Universal QUICK-Clone cDNA II (Clontech) for a template cDNA and then cloned into a pET-41 Ek/LIC vector (Novagen). Recombinant proteins of Rab5a-His12, Rab5aΔHVR-His12, Rab4a-His12, Rab4aΔHVR-His12, and untagged Rab4aΔHVR (**Figure 1**) were expressed in *Escherichia coli* BL21(DE3) cells (Novagen) harboring the pET-41-based vectors constructed. After inducing protein expression by adding IPTG (0.2 mM final, 37°C, 3 hours), cultured cells were harvested, resuspended with the RB150 buffer (20 mM Hepes-NaOH, pH 7.4, 150 mM NaCl, 10% glycerol) containing 0.1 mM GTP, 5 mM MgCl2, 1 mM DTT, 1 mM PMSF, and 1 μg/ml pepstatin A, freeze-thawed in liquid nitrogen and a water bath at 30°C, lysed by sonication, and ultracentrifuged with a 70 Ti rotor (Beckman Coulter; 50,000 rpm, 75 min, 4°C). Supernatants after ultracentrifugation were mixed with COSMOGEL GST-Accept beads (Nacalai Tesque) and incubated with gentle agitation (4°C, 2 hours). The protein-bound beads were washed four times in RB150 containing 5 mM MgCl2 and 1 mM DTT, resuspended in the same buffer, supplemented with human rhinovirus 3C protease (8 units/ml; Novagen), and incubated without agitation (4°C, 16 hours). Purified Rab proteins, which had only three extra residues (Gly-Pro-Gly) at the N-terminus, were eluted from the beads after proteolytic cleavage and analyzed by SDS-PAGE and CBB staining (**Figure 1**). Concentrations of purified Rab proteins were determined using Protein Assay CBB Solution (Nacalai Tesque) and BSA as a standard protein.

**Figure 1.**
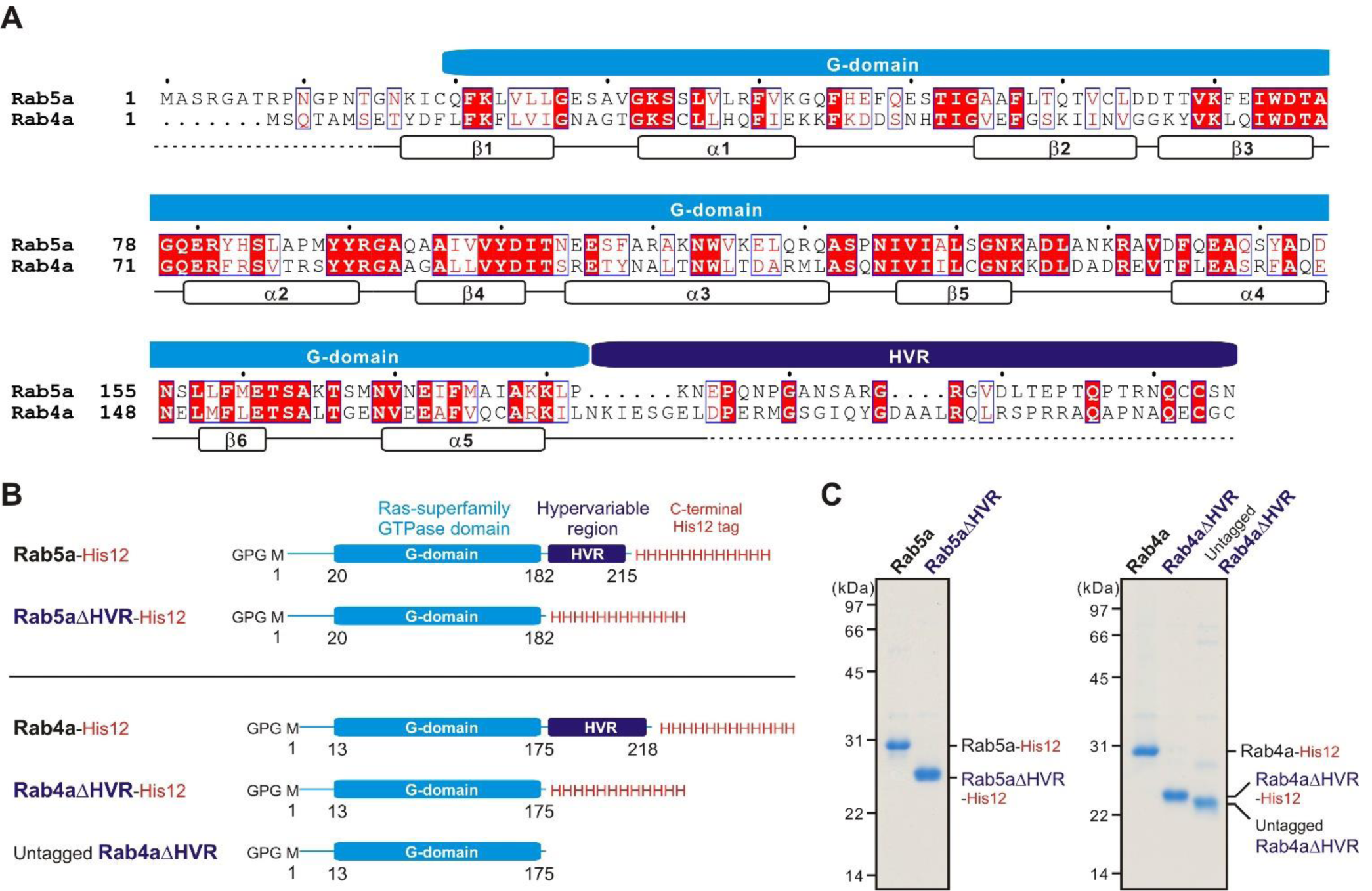
Human endosomal Rab-family small GTPases used in the current reconstitution studies. (**A**) Sequence alignment of human Rab5a and Rab4a proteins. Amino acid sequences of the two endosomal Rab isoforms were obtained from UniProtKB (https://www.uniprot.org/), aligned using ClustalW (https://www.genome.jp/tools-bin/clustalw), and rendered by ESPript 3.0 (http://espript.ibcp.fr/ESPript/ESPript/). Identical and similar amino acid residues in the sequence alignment are highlighted in red boxes and in red characters, respectively. Sequence regions corresponding to the conserved Ras superfamily GTPase domain (G-domain) and the C-terminal hypervariable region (HVR) are indicated on the top of the alignment. Secondary structures determined by the crystal structure of human Rab5a containing the residues 15-184 (PDB code, 1N6H), including five α-helices (α1-α5) and six β-strands (β1-β6), are indicated at the bottom of the alignment. (**B**) Schematic representation of recombinant proteins of human endosomal Rabs used in the current studies, which include the C-terminally His12-tagged forms of full-length Rab5a (Rab5a-His12), HVR-deleted mutant Rab5a (Rab5aΔHVR-His12), full-length Rab4a (Rab4a-His12), and HVR-deleted mutant Rab4a (Rab4aΔHVR-His12), and the untagged form of HVR-deleted mutant Rab4a (untagged Rab4aΔHVR). All of the wild-type and mutant Rab proteins have only three extra residues (Gly-Pro-Gly) at the N-terminus after purification. (**C**) Coomassie blue-stained gels of purified Rab5a-His12, Rab5aΔHVR-His12, Rab4a-His12, Rab4aΔHVR-His12, and untagged Rab4aΔHVR proteins, which were tested in the reconstituted liposome tethering assays in Figures 2-7.

### Liposome preparation

For preparing synthetic protein-free liposomes, all of the non-fluorescent lipids used, including POPC (1-palmitoyl-2-oleoyl-phosphatidylcholine), POPE (1-palmitoyl-2-oleoyl-phosphatidylethanol), bovine liver PI (phosphatidylinositol), POPS (1-palmitoyl-2-oleoyl-phosphatidylserine), ovine wool cholesterol, and DOGS-NTA (1,2-dioleoyl-sn-glycero-3-{[N-(5-amino-1-carboxypentyl) iminodiacetic acid]-succinyl}), were purchased from Avanti Polar Lipids, and the two fluorescence-labeled lipids used, Rh-PE (rhodamine-PE) and FL-PE (fluorescein-PE), were from Molecular Probes. Lipids were mixed in chloroform with the lipid compositions of 41% (mol/mol) POPC, 17% POPE, 10% liver PI, 5% POPS, 20% ovine cholesterol, 6% DOGS-NTA, and 1% Rh-PE or FL-PE. After evaporating chloroform with a stream of nitrogen gas, dried lipid mixes were resuspended in RB150 containing 5 mM MgCl2 and 1 mM DTT by vortexing (final 8 mM total lipids), incubated with agitation (37°C, 1 hour), freeze-thawed in liquid nitrogen and a water bath at 30°C, and extruded 25 times through polycarbonate filters (pore diameters, 200 nm; Avanti Polar Lipids) in a mini-extruder (Avanti Polar Lipids) preheated at 40°C. The liposome solutions prepared were stored at 4°C and used within a week for all of the current reconstitution experiments. Size distributions of the extruded liposomes were measured by dynamic light scattering (DLS) using a DynaPro NanoStar DLS instrument (Wyatt Technology) (**Figure S1**).

### Liposome turbidity assay

To quantitatively evaluate the intrinsic capacities of human Rab-family small GTPases to physically tether two distinct lipid membranes, turbidity changes of liposome solutions in the presence of purified Rab proteins were monitored by measuring optical density at 400 nm, as described (Tamura and Mima, 2014; Inoshita and Mima, 2017; Mima, 2018; Segawa et al., 2019). Liposome solutions (200-nm diameter; 1 mM total lipids in final) and purified Rab-His12 proteins (final 0.2-10 μM), which had been preincubated separately at 30°C for 5 min, were mixed in RB150 containing 5 mM MgCl2 and 1 mM DTT, transferred to a 10-mm path-length cuvette (105.201-QS, Hellma Analytics), and immediately subjected to measurement of the optical density changes at 400 nm (ΔOD400) in a DU720 spectrophotometer (Beckman Coulter) for 5 min with 10-sec intervals at room temperature. The ΔOD400 data obtained from the kinetic turbidity assays were analyzed by curve fitting using the ImageJ2 software (National Institutes of Health) and the logistic function formula, y = a/(1+b*exp(-c*x)), where y and x correspond to the ΔOD400 value and the time (min), respectively (Segawa et al., 2019; Taniguchi et al., 2020). The maximum capacities of Rab-mediated liposome tethering were defined as the theoretical maximum ΔOD400 values of the fitted sigmoidal curves at *t* = ∞ and thus calculated as “a” from the logistic formula above. In addition, the initial velocities of liposome tethering were defined as the maximum slopes of the fitted curves and calculated as “c*a/4” from the formula above. Means and standard deviations of the tethering capacities and velocities were determined from three independent experiments. The turbidity data were statistically evaluated using two-way ANOVA in SigmaPlot 11 (Systat Software). All of the kinetic plots shown in the turbidity assays were obtained from one experiment and were typical of those from more than three independent experiments.

### Fluorescence microscopy

Fluorescence microscopy-based imaging assays for Rab-mediated liposome tethering were performed using a LUNA-FL automated fluorescence cell counter (Logos Biosystems), as described (Segawa et al., 2019; Taniguchi et al., 2020). Liposomes bearing fluorescence-labeled Rh-PE or FL-PE lipids (200-nm diameter; final 2 mM total lipids) and Rab-His12 proteins (final 0.5-8 μM), which had been separately preincubated (30°C, 10 min), were mixed in RB150 with 5 mM MgCl2 and 1 mM DTT, incubated without agitation (30°C, 2 hours), and then applied to a LUNA cell-counting slide (L12001, Logos Biosystems; 15 μl per well). Bright field images, Rh-fluorescence images, and FL-fluorescence images of the Rab-mediated liposome tethering reactions in the slides were obtained and processed by the LUNA-FL cell counter. Particle sizes of Rab-dependent liposome clusters observed in the fluorescence images were analyzed using the ImageJ2 software with setting the lower intensity threshold level to 150, the upper intensity threshold level to 255, and the minimum particle size to 10 pixel^2^ which corresponds to approximately 10 μm^2^ (Segawa et al., 2019; Taniguchi et al., 2020).

## Results

Rab-family small GTPases constitute the largest branch of the Ras superfamily, which includes 11 Ypt/Rab proteins in budding yeast and more than 60 Rab isoforms in humans (Rojas et al., 2012). In general, Rab proteins from all eukaryotes are a small monomeric protein of approximately 25 kDa and are comprised of the Ras superfamily small GTPase domain (G-domain; 160-170 residues), which can specifically associate with the cognate interacting proteins (or protein complexes) called “Rab effectors” in a GTP-dependent manner to mediate the multiple steps of intracellular membrane trafficking as a molecular switch (Zerial and McBride, 2001; Stenmark, 2009; Hutagalung and Novick, 2011), and also two other non-conserved regions adjacent to the conserved globular G-domain (Khan and Ménétrey, 2013; Pylypenko et al., 2018; Mima, 2018), which include the flexible N-terminal segment (5-30 residues) and the unstructured C-terminal HVR domain (20-50 residues) that has been studied on its contributions to intracellular localization of Rab GTPases (Ali et al., 2004; Li et al., 2014) and recently reported to be involved in selective interaction with the guanine nucleotide exchange factors (Thomas et al., 2019). Notably, in addition to these conventional structural and functional features of Rab-family GTPases (Zerial and McBride, 2001; Stenmark, 2009; Hutagalung and Novick, 2011), recent reconstitution studies have revealed their novel molecular functions to directly and physically tether lipid membranes by themselves (Lo et al., 2012; Tamura and Mima, 2014; Inoshita and Mima, 2017; Mima, 2018; Segawa et al., 2019). Our comprehensive experiments for 11 representative human Rab isoforms (Rab1a, −3a, −4a, −5a, - 6a, −7a, −9a, −11a, −14, −27a, and −33b) demonstrated that the intrinsic tethering capacities are highly conserved among all of the Rabs tested, except for Rab27a, and are achieved exclusively through *trans*-assembly between membrane-anchored Rab proteins in homotypic and heterotypic Rab combinations (Inoshita and Mima, 2017; Segawa et al., 2019). Here, based on the earlier findings above by an in vitro reconstitution approach, we further explored molecular mechanisms by which Rab proteins confer efficiency and specificity of their tethering activities, particularly focusing on the roles of the C-terminal HVR flexible linkers that connect the globular G-domains to membrane surfaces.

### Deletion of the HVR linkers enhances the intrinsic tethering potency of human Rab proteins

For thoroughly comparing the intrinsic tethering activities of the full-length, wild-type form and the HVR-deleted mutant form of Rab-family proteins, we selected the two human Rab isoforms, Rab5a and Rab4a, as a typical model among over 60 Rab members in human cells (**Figure 1**). These two Rab isoforms, which are both principally localized at the cytoplasmic face of early endosomal membranes (Zerial and McBride, 2001; Stenmark, 2009; Hutagalung and Novick, 2011), exhibit more than 40% sequence identity with their G-domains, but they have little or no conserved sequence or motif in their HVR linker domains (**Figure 1A**). It is also noteworthy that early endosomal Rab5a and Rab4a proteins were found to be typical of a highly-potent membrane tether and an inefficient membrane tether, respectively (Inoshita and Mima, 2017; Segawa et al., 2019).

As tested in our prior works on Rab-mediated tethering, recombinant proteins of full-length Rab5a, Rab4a, and their HVR-deleted mutants (denoted as Rab5aΔHVR and Rab4aΔHVR) were purified in the C-terminally-modified forms with an artificial His12 tag (**Figure 1B, C**), which allows purified Rab proteins to stably associate with lipid bilayers of synthetic liposomes bearing a DOGS-NTA lipid (**Figure 2A**), mimicking membrane attachment of native Rab proteins via an isoprenyl lipid anchor at the C-terminus (Mima, 2018). Liposomes used in the current reconstitution systems were prepared by an extrusion method with a 200-nm pore-size filter (**Figure 2A**), yielding the curvature of lipid bilayers roughly similar to that of early endosomal membranes in mammalian cells, which were shown to be approximately 100-500 nm in diameter (Klumperman and Raposo, 2014). Regarding the lipid composition, the extruded 200-nm liposomes bore five major lipid species, including PC, PE, PI, PS, and cholesterol (**Figure 2A**), which primarily compose organelle membranes in mammals (van Meer et al., 2008; Vance, 2015; Yang et al., 2018).

**Figure 2.**
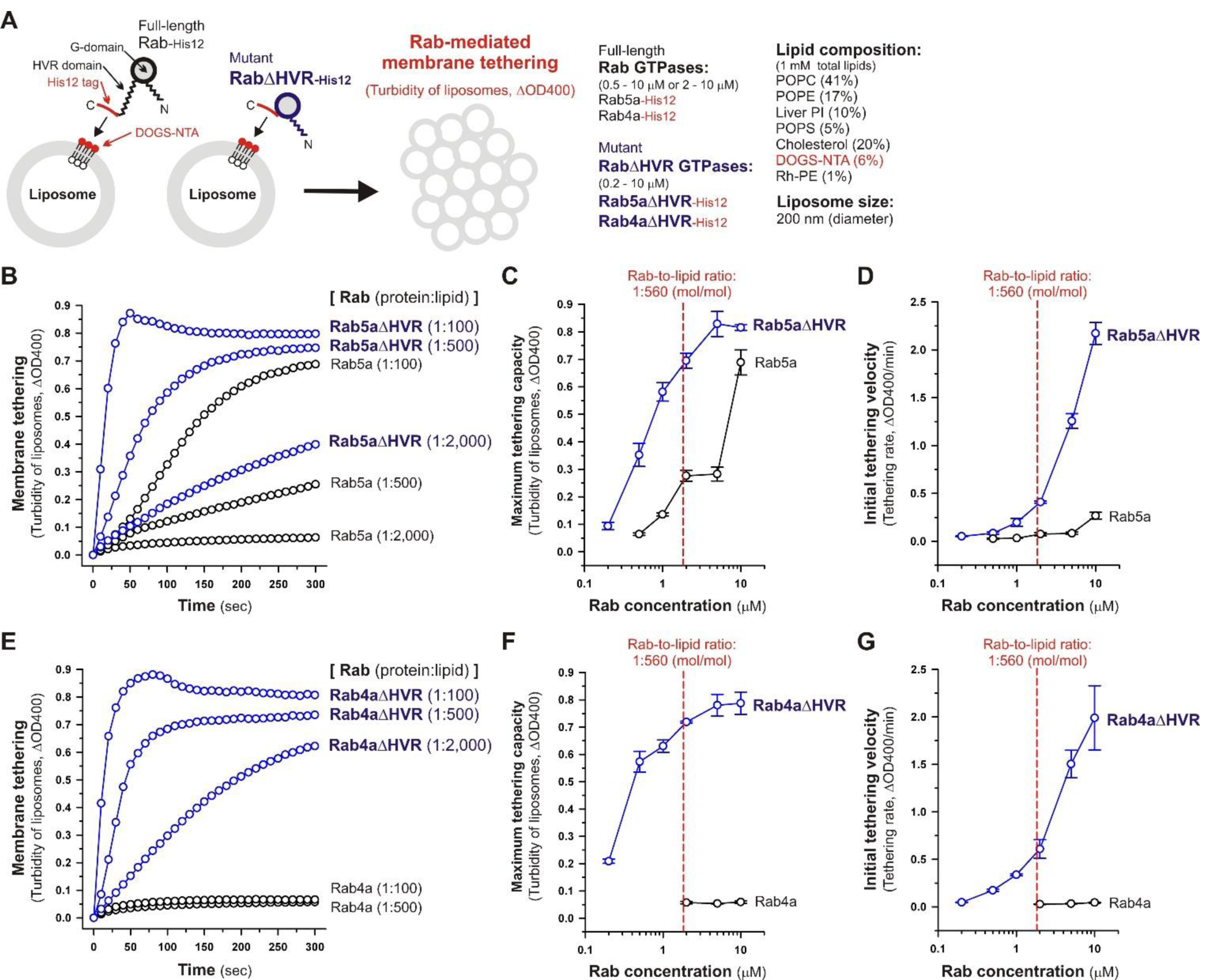
Deletion of the flexible HVR linkers enhances membrane tethering potency of human Rab5a and Rab4a small GTPases. (**A**) Schematic representation of membrane tethering assays for human endosomal Rab5a and Rab4a in a chemically-defined reconstitution system. (**B-D**) Kinetic liposome turbidity assays for Rab5a-mediated membrane tethering. Purified Rab5a-His12 (final 0.5-10 μM) and Rab5aΔHVR-His12 (final 0.2-10 μM) were mixed with DOGS-NTA-bearing synthetic liposomes (200-nm diameter; final 1 mM total lipids) and immediately assayed for the turbidity changes by measuring the optical density at 400 nm (ΔOD400) for 5 min (**B**). The kinetic turbidity data were further analyzed by a sigmoidal curve fitting method to determine the maximum tethering capacities (**C**) and the initial tethering velocities (**D**). (**E-G**) Kinetic liposome turbidity assays for Rab4a-mediated membrane tethering. Purified Rab4a-His12 (final 2-10 μM) and Rab4aΔHVR-His12 (final 0.2-10 μM) were mixed with DOGS-NTA-bearing liposomes and assayed for the turbidity changes (**E**), as in (**B**). The kinetic turbidity data were analyzed by curve fitting, thereby determining the maximum capacities (**F**) and the initial velocities (**G**), as in (**C, D**). The Rab protein concentration at the physiological Rab-to-lipid molar ratio of 1:560 (mol/mol) is indicated as a red dashed line (**C-D, F-G**). Error bars, SD.

Under the physiologically-relevant conditions, the kinetic turbidity assays for liposome tethering were employed with full-length and HVR-deleted Rab5a (**Figure 2B-D**) and Rab4a (**Figure 2E-G**) to analyze the tethering capacities at a broad range of the Rab protein-to-lipid ratios, from 1:5,000 to 1:100 (mol/mol). Strikingly, the current liposome tethering assays uncovered that both Rab5a and Rab4a isoforms can greatly stimulate the intrinsic tethering activities by removal of their HVR linker domains that are located between the G-domains and C-terminal membrane anchors (**Figure 2B, E**). In the case of Rab5a, which had been reported to be the most tethering-competent isoform among human Rab-family proteins tested (Tamura and Mima, 2014; Inoshita and Mima, 2017; Segawa et al., 2019), Rab5aΔHVR mutant exhibited over 5-fold higher maximum tethering capacities and up to 15-fold higher initial tethering velocities than those values for wild-type Rab5a (**Figure 2C, D**). This is basically consistent with modest increase in the tethering activity of the yeast Rab5 ortholog, Vps21p, by truncation of its C-terminal linker (Lo et al., 2012). Likewise, the HVR-deleted mutant of Rab4a (Rab4aΔHVR) exhibited the very high tethering potency comparable to that of Rab5aΔHVR (**Figure 2B, E**), yielding more than 10-fold higher maximum tethering capacities (**Figure 2F**) and over 40-fold higher initial tethering velocities (**Figure 2G**) compared to full-length Rab4a that showed little tethering activities under the current conditions with the 200-nm liposomes (**Figure 2E-G**).

Considering the physiological Rab-to-lipid molar ratio (1:560, mol/mol; **Figure 2C, D, F, G**, red dashed lines), which was calculated as described (Inoshita and Mima, 2017; Segawa et al., 2019) using the average copy number of Rab proteins in synaptic vesicles (25 Rab molecules per vesicle; Takamori et al., 2006), the mean diameter of synaptic vesicles (42 nm; Takamori et al., 2006), the typical thickness of biological membranes (4 nm; Nagle and Tristram-Nagle, 2000), and the average surface area of phospholipid headgroups (0.65 nm^2^; Nagle and Tristram-Nagle, 2000), kinetic data of the reconstituted tethering assays further demonstrated that the HVR-deleted forms of both Rab5a and Rab4a have the intrinsic potency to drive rapid and efficient tethering of liposomal membranes at the physiologically-relevant Rab protein densities on membrane surfaces and even at much lower Rab densities, such as the Rab-to-lipid molar ratio of 1:2,000 (mol/mol; **Figure 2B-G**). Although we used the protein stoichiometry of synaptic vesicles as a reasonable model for calculating the physiological Rab protein density on the membrane (Takamori et al., 2006), it is possible that the Rab densities on other membrane compartments such as early endosomes are variable and different from the current estimations. Regarding other putative membrane tethers in a reconstituted system, their tethering activities have been examined at the protein-to-lipid ratios similar to those tested in the present experiments (1:100-1:5,000, mol/mol; **Figure 2**), which include the typical ratios of 1:400 for golgin GMAP-210 (Drin et al., 2008), 1:330 for Vps21p (Lo et al., 2012), 1:2,000 for HOPS (Ho and Stroupe, 2015; Ho and Stroupe, 2016), 1:800 for Atg8p (Nair et al., 2011), and 1:500 for human Atg8 orthologs (Taniguchi et al., 2020). It should also be noted that, assuming that Rab molecules are a spherical 25-kDa protein with a radius of 2.0 nm (Erickson, 2009), membrane-bound Rab proteins occupy only 1.9% of the outer surface areas of the 200-nm liposomes when tested at the 1:2,000 Rab-to-lipid ratio. Thus, this reflects that membrane tethering driven by HVR-deleted Rabs is a highly robust and specific biochemical reaction in the physiological context and is quite unlikely to be caused by non-selective protein-protein or protein-lipid interactions on membrane surfaces.

Microscopic observations of fluorescence-labeled liposome clusters induced by membrane-anchored Rab proteins, as an alternative assay for Rab-mediated membrane tethering, provided further experimental evidence of the high tethering potency of the HVR-deleted forms of human Rab-family proteins (**Figure 3**). To comprehensively evaluate the intrinsic capacities of full-length and HVR-deleted Rab5a and Rab4a proteins to induce the formation of massive clusters of fluorescent Rh-PE-bearing liposomes (**Figure 3A**), the wide-field Rh-fluorescence images of the Rab-mediated liposome tethering reactions, which had been incubated (2 hours) with the Rh-PE-bearing 200-nm liposomes and Rab5a (**Figure 3B**), Rab5aΔHVR (**Figure 3C**), Rab4a (**Figure 3G**), or Rab4aΔHVR (**Figure 3H**), were acquired using a LUNA-FL fluorescence cell counter and a LUNA cell-counting slide. The current imaging assays for liposome tethering allowed us to simultaneously observe large numbers of the Rab-induced liposome clusters formed in a defined volume (length × width × height = 2,500 × 2,000 × 100 μm), thereby unbiasedly and quantitatively measuring their particle numbers (**Figure 3D, I**), average particle sizes (**Figure 3E, J**), and total particle areas (**Figure 3F, K**). Consistent with the results from the kinetic turbidity assays (**Figure 2B-G**), both Rab5aΔHVR and Rab4aΔHVR mutant proteins were able to trigger highly efficient liposome tethering in the imaging assays, yielding more than 400 detectable particles of Rh-labeled liposome clusters (**Figure 3D, I**) with the average sizes above 700 μm^2^ (**Figure 3E, J**) in the Rh-fluorescence images obtained (**Figure 3C-F, Figure 3H-K**). Notably, when assayed at the Rab-to-lipid molar ratios of 1:2,000 (final 1 μM Rabs and 2 mM lipids), these HVR-deleted Rab proteins yielded the total areas of liposome clusters ranging from 300,000 to 500,000 μm^2^ (**Figure 3F, K**), whereas full-length Rab5a and Rab4a proteins were almost incompetent to initiate efficient liposome tethering under the conditions (**Figure 3B, G**), giving only 1,000 to 2,000 μm^2^ for the total particle areas (**Figure 3F, K**). In addition to these findings on very high tethering potency of the HVR-deleted mutants of Rab5a and Rab4a (**Figures 2, 3**), it should also be noted that, although full-length Rab5a exhibited its adequate tethering potency at the Rab-to-lipid molar ratios of 1:1,000 and 1:500 in the microscopic assays, it turned out to be totally inactive at the higher Rab-to-lipid ratio of 1:250 (**Figure 3B, D-F**). This appears to be puzzling but perhaps reflects that full-length Rab5a is prone to assemble into a homo-dimeric complex in the *cis*-configuration at such high Rab densities, preventing the protein assemblies in *trans* between two opposing membranes, while the HVR-deleted proteins still rather assemble into the *trans*-complexes under the same conditions.

**Figure 3.**
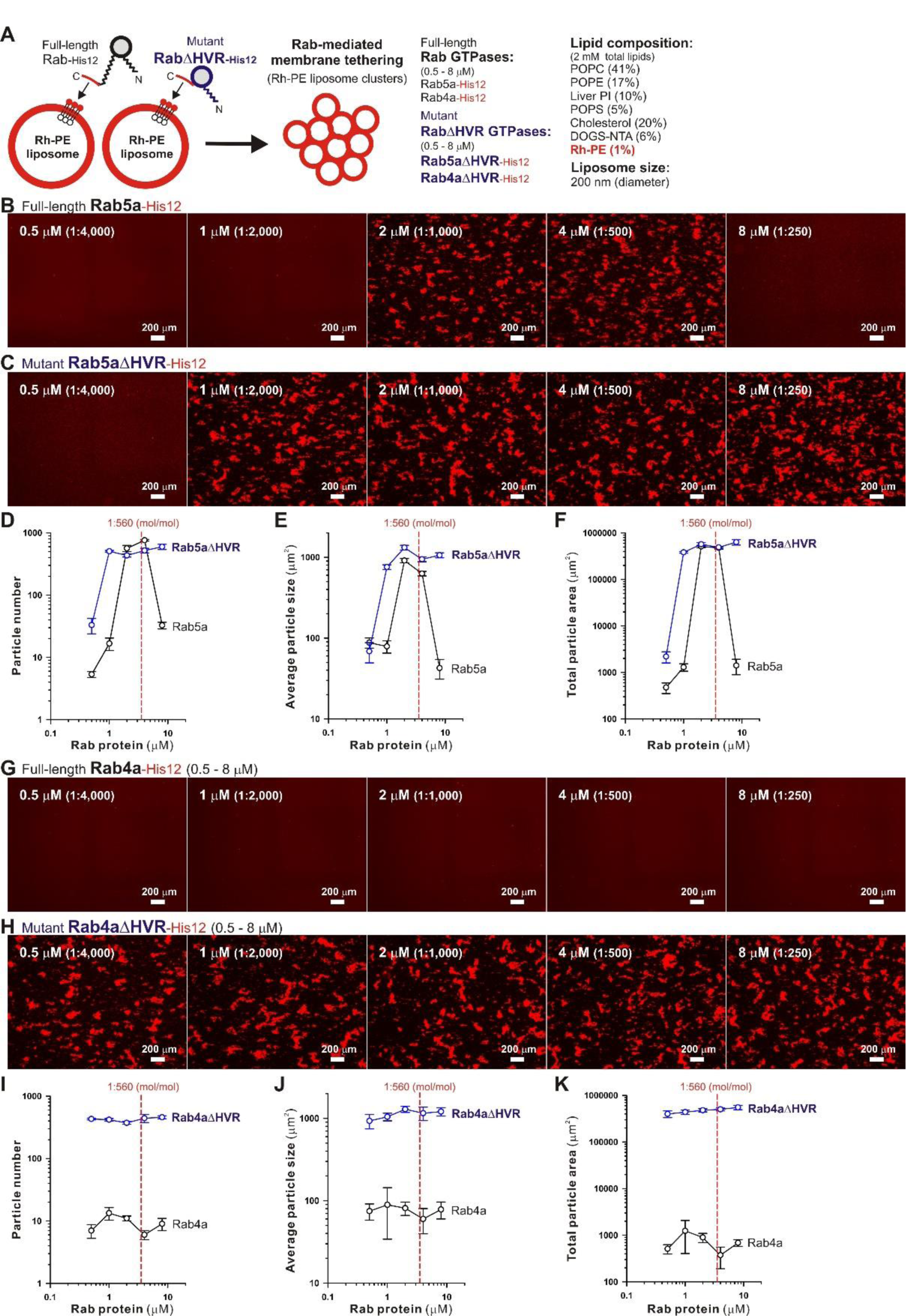
Fluorescence microscopic analysis of membrane tethering driven by HVR-deleted Rab proteins. (**A**) Schematic representation of fluorescence microscopy-based assays for membrane tethering driven by full-length and HVR-deleted forms of endosomal Rab5a and Rab4a. (**B-K**) Purified Rab5a-His12 (**B**), Rab5aDHVR-His12 (**C**), Rab4a-His12 (**G**), and Rab4aDHVR-His12 (**H**) proteins (final 0.5-8 μM) were mixed with fluorescence-labeled liposomes bearing Rh-PE and DOGS-NTA (200-nm diameter; final 2 mM total lipids), incubated (30°C, 2 hours), and subjected to fluorescence microscopy. Particle sizes of Rab-induced liposome clusters in the rhodamine fluorescence images obtained (**B-C, G-H**) were measured using the ImageJ2 software, yielding means and SD values of the particle numbers (**D, I**), average particle sizes (**E, J**), and total particle areas (**F, K**), which were determined from three independent images of the Rab-mediated liposome tethering reactions. Scale bars, 200 μm. Error bars, SD.

### Requirement of *trans*-assembly of HVR-deleted Rab proteins in reversible membrane tethering reactions

Next, we further employed the microscopic imaging assays for liposome clustering to ask whether membrane tethering mediated by HVR-deleted Rab mutant proteins is a non-fusogenic, reversible tethering reaction (**Figure 4**) and also whether *trans*-assembly between membrane-anchored Rab proteins is certainly required for the tethering events driven by HVR-deleted Rabs (**Figure 5**), as previously established for full-length wild-type Rabs (Tamura and Mima, 2014; Inoshita and Mima, 2017; Segawa et al., 2019). To test reversibility of the tethering reactions with Rab4aΔHVR-His12 proteins and DOGS-NTA-bearing fluorescent liposomes (**Figure 4**), large clusters of Rab4aΔHVR-anchored liposomes, which had been pre-formed during the first incubation (1 hour), were supplemented with imidazole (250 mM), which acts as a competitive inhibitor to block the membrane association of Rab-His12 proteins, or with the buffer control, further incubated (1 hour), and then subjected to fluorescence microscopy (**Figure 4A**). Obviously, although a number of huge Rh-labeled liposome clusters were still present in the reaction incubated with the buffer control (the total particle area of 410,000 μm^2^; **Figure 4B**), addition of imidazole caused untethering of liposome clusters and completely abolished detectable Rh-labeled particles (the total particle area of 2,000 μm^2^; **Figure 4C**). These results demonstrated reversible membrane tethering mediated by HVR-deleted Rab proteins that can be strictly controlled by the Rab attachment and detachment cycles on membrane surfaces, consistent with the earlier experimental evidence for the reversibility of tethering between yeast vacuoles (Mayer and Wickner, 1997; Ungermann et al., 1998) and tethering between full-length Rab-anchored liposomes (Tamura and Mima, 2014).

**Figure 4.**
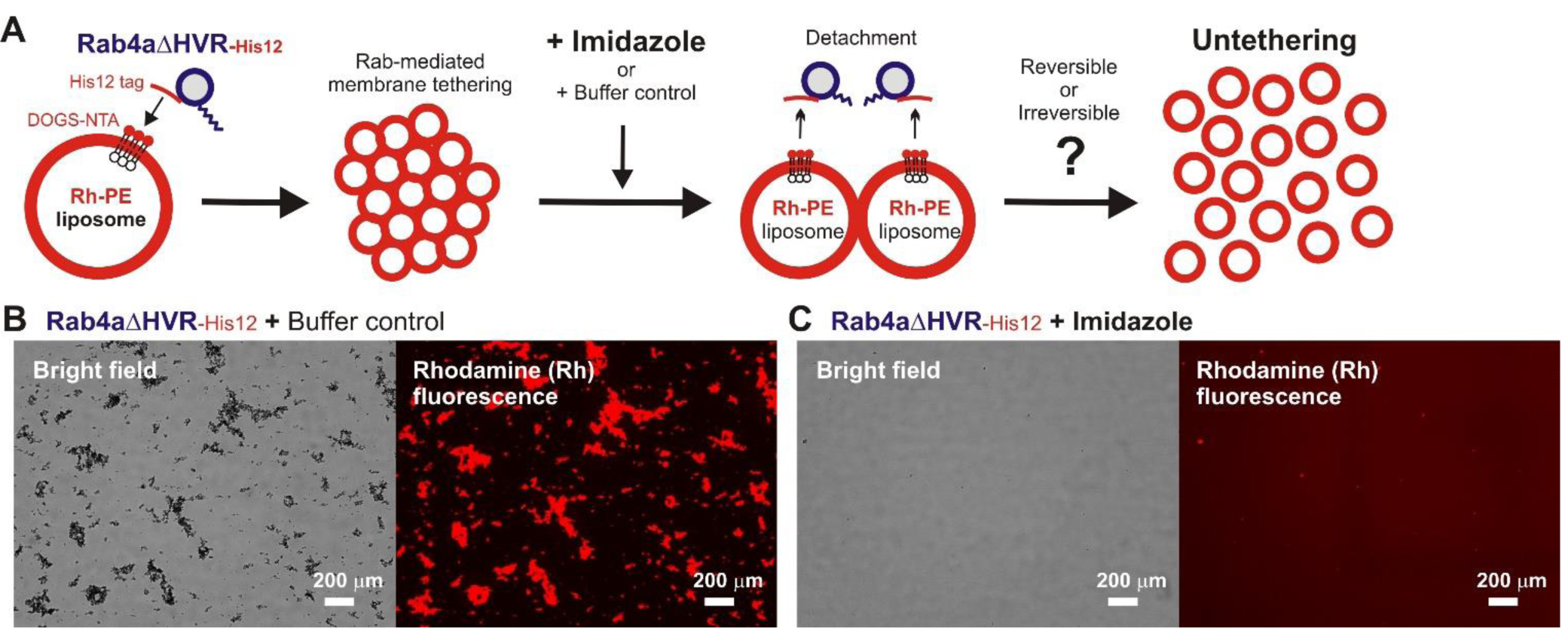
HVR-deleted Rab4a drives a non-fusogenic, reversible membrane tethering reaction. (**A**) Schematic representation of fluorescence microscopy-based assays testing the reversibility of reconstituted membrane tethering mediated by HVR-deleted mutant Rab4a. (**B-C**) Purified Rab4aΔHVR-His12 (final 1 μM) and fluorescence-labeled liposomes bearing Rh-PE and DOGS-NTA (200-nm diameter; final 2 mM total lipids) were mixed and incubated (30°C, 1 hour) to induce the formation of Rab4aΔHVR-mediated liposome clusters. After the first incubation, the liposome tethering reactions were supplemented with the buffer control (**B**) or imidazole (final 250 mM) which causes the dissociation of Rab4aΔHVR-His12 proteins from DOGS-NTA-bearing liposomes (**C**), further incubated (30°C, 1 hour), and analyzed by fluorescence microscopy to obtain the bright field images and rhodamine fluorescence images. Particle sizes of Rab-induced liposome clusters in the rhodamine fluorescence images (**B, C**) were measured using the ImageJ2 software as in Figure 3, yielding the total particle areas of 360,000 ± 48,000 μm^2^ (**B**) and 1,700 ± 790 μm^2^ (**C**), which were determined from three independent fluorescence images of the tethering reactions. Scale bars, 200 μm.

**Figure 5.**
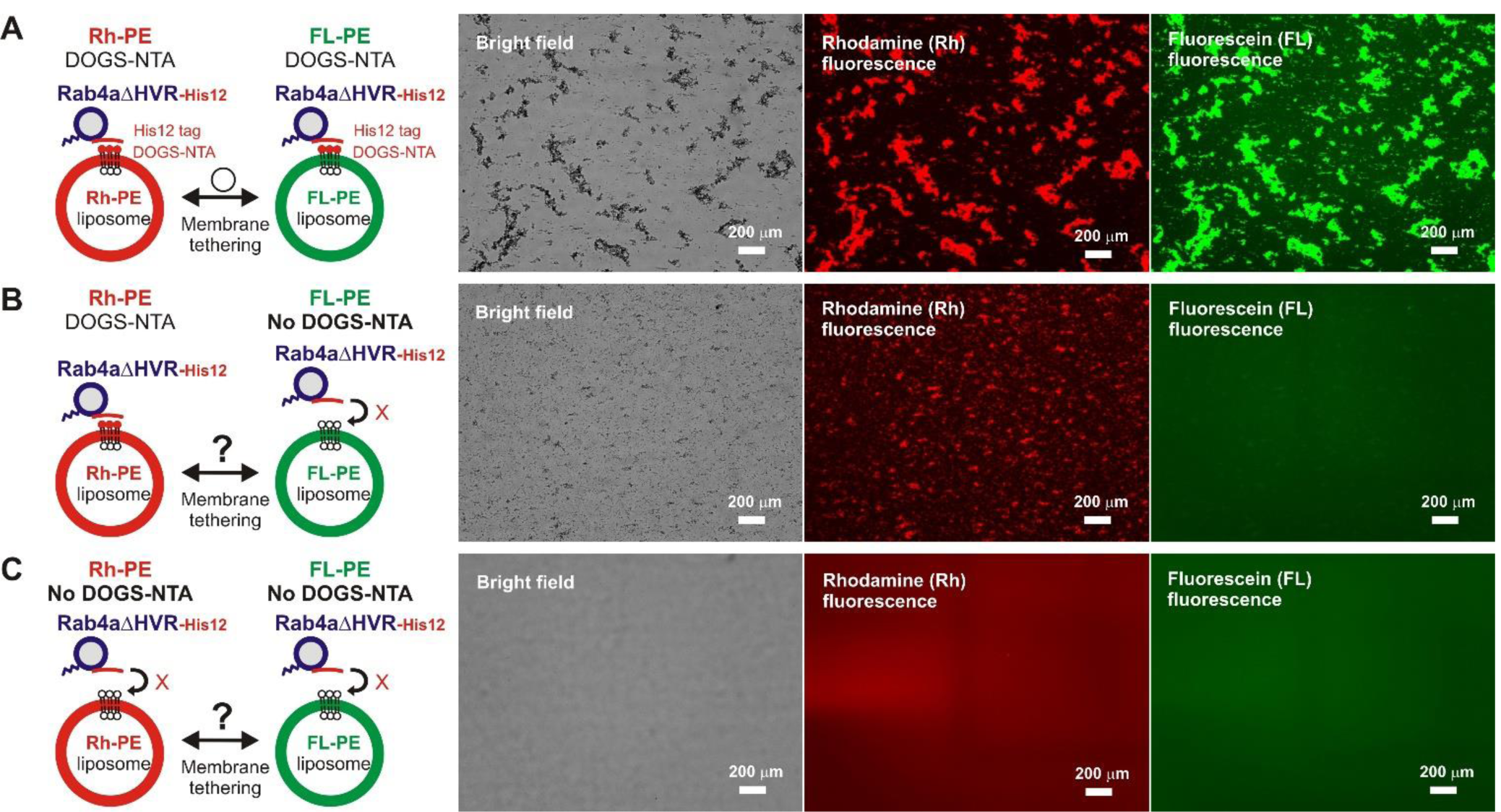
*Trans*-assembly between membrane-anchored Rab proteins is required for membrane tethering mediated by HVR-deleted Rab4a. (**A-C**) Fluorescence microscopy analysis of reconstituted membrane tethering driven by *trans*-assembly of membrane-anchored Rab4aΔHVR proteins. Purified Rab4aΔHVR-His12 (final 2 μM) and two types of fluorescence-labeled liposomes, Rh-PE-bearing liposomes and FL-PE-bearing liposomes (200-nm diameter; final 1 mM total lipids for each), were mixed, incubated (30°C, 1 hour), and subjected to fluorescence microscopy, obtaining the bright field images (left panels), rhodamine fluorescence images (middle panels), and fluorescein fluorescence images (right panels). DOGS-NTA lipids for membrane anchoring of Rab4aΔHVR-His12 proteins were present in both the Rh-PE and FL-PE liposomes (**A**), only in the Rh-PE liposomes (**B**), or not present in either of the liposomes (**C**). Particle sizes of Rab-induced liposome clusters in the rhodamine fluorescence images (**A-C**) were measured using the ImageJ2 software as in Figure 3, yielding the total particle areas of 450,000 ± 42,000 μm^2^ (**A**), 160,000 ± 39,000 μm^2^ (**B**), and 170 ± 80 μm^2^ (**C**), which were determined from three independent fluorescence images of the tethering reactions. Scale bars, 200 μm.

For exploring the requirement of Rab-Rab assemblies in *trans* during HVR-deleted Rab-mediated tethering, we performed the same microscopic assays but using two types of the fluorescence-labeled DOGS-NTA-liposomes which bore either Rh-PE or FL-PE (**Figure 5**). As expected, massive liposome clusters induced by Rab4aΔHVR-His12 proteins contained both of the Rh-PE-liposomes (**Figure 5A**, middle panel) and FL-PE-liposomes (**Figure 5A**, right panel), yielding 490,000 μm^2^ for the total particle area of the Rh-labeled clusters that entirely overlapped with the FL-labeled clusters. However, by omitting DOGS-NTA from the FL-PE-liposomes, HVR-deleted Rab4a no longer had the capacity to form large FL-labeled clusters (**Figure 5B**, right panel), only inducing the Rh-labeled liposome clusters (**Figure 5B**, middle panel; the total particle area of 140,000 μm^2^). Moreover, the omission of DOGS-NTA from both fluorescence-labeled liposomes completely abrogated the intrinsic tethering activity of Rab4aΔHVR-His12 proteins (**Figure 5C**). These data clearly establish that highly efficient membrane tethering driven by HVR-deleted Rab proteins requires their membrane-bound forms on both opposing membranes, thus reflecting the need for *trans*-Rab-Rab assemblies to bridge two distinct lipid bilayers destined to be tethered.

Specific *trans*-assembly of the membrane-anchored form of HVR-deleted Rab proteins during reconstituted membrane tethering was further investigated by employing liposome turbidity assays in the presence of untagged Rab4aΔHVR that lacks a His12 tag at the C-terminus (**Figure 6**). Before initiating Rab-dependent liposome tethering by mixing with DOGS-NTA-bearing liposomes, Rab4aΔHVR-His12 was pre-incubated (10 min) with an up to 10-fold molar excess of untagged Rab4aΔHVR that potentially inhibits the *trans*-interactions between membrane-bound Rab4aΔHVR-His12 proteins (**Figure 6A**). Nevertheless, the presence of excess untagged Rab4aΔHVR proteins in solution had little effect on the tethering capacity of membrane-bound Rab4aΔHVR-His12 proteins (**Figure 6B**, see the ΔOD400 values at 300 sec), even though the initial rates of tethering were slightly reduced by the addition of untagged Rab4aΔHVR (**Figure 6B**), suggesting the weak interactions between untagged Rab4aΔHVR and Rab4aΔHVR-His12 that prevent rapid Rab-Rab assemblies in *trans*. These results lead us to conclude that rapid and efficient membrane tethering driven by HVR-deleted Rab proteins are achieved by highly selective *trans*-Rab-Rab interactions, distinguishing membrane-bound Rabs from the membrane-unbound soluble forms.

**Figure 6.**
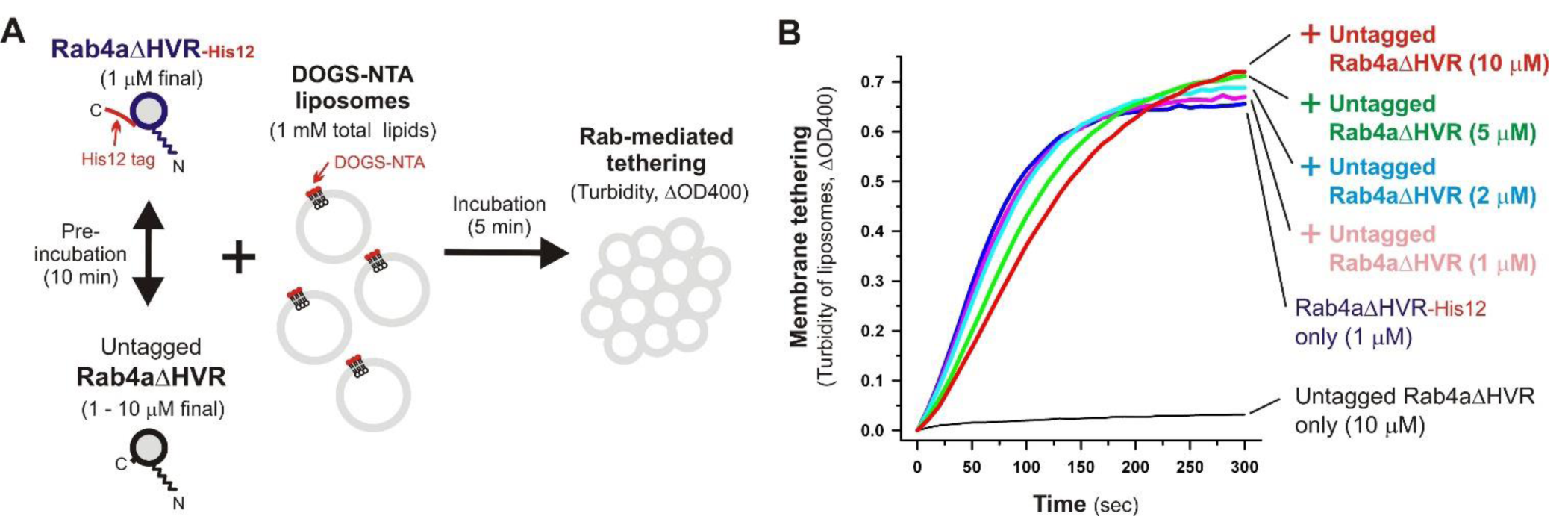
Selective *trans*-assembly of HVR-deleted Rab4a proteins in reconstituted membrane tethering. (**A**) Schematic representation of liposome turbidity assays in (**B**), testing the inhibitory effect of soluble untagged Rab4aΔHVR proteins lacking a His12-tag membrane anchor on membrane tethering reactions driven by *trans*-assembly of membrane-anchored Rab4aΔHVR-His12 proteins. (**B**) Rab4aΔHVR-His12 proteins (final 1 μM) were mixed with equimolar or excess amounts of untagged Rab4aΔHVR proteins (final 1-10 μM) and incubated at 30°C for 10 min. After pre-incubation, protein mixtures of Rab4aΔHVR-His12 and untagged Rab4aΔHVR were added to the suspensions containing DOGS-NTA-bearing liposomes (200-nm diameter; final 1 mM total lipids), immediately followed by monitoring the turbidity changes by measuring the optical density at 400 nm (ΔOD400) for 5 min, as in Figure 2.

### Effects of membrane lipids on membrane tethering functions of full-length and HVR-deleted Rab proteins

The current reconstitution experiments with the full-length and HVR-deleted forms of human Rab-family small GTPases established that deletion of the C-terminal HVR domains can enhance the intrinsic tethering potency of human Rab proteins (e.g., endosomal Rab5a and Rab4a) to trigger reversible membrane tethering mediated by highly selective Rab-Rab protein assemblies in *trans* (**Figures 2-6**). Since the HVR domains are a 20-50 residue long flexible linker that connects the G-domain to a membrane anchor at the C-terminus (Khan and Ménétrey, 2013; Mima, 2018; **Figure 1, Figure 2A**), removal of the HVR linkers greatly shortens the distance between the globular G-domains and membranes when anchored to lipid bilayers, allowing the G-domains to be placed in close contact with membrane surfaces. Thus, the present data shown in **Figures 2-6** faithfully reflect that the close membrane attachment of the Rab G-domains is an essential process to promote rapid and efficient *trans*-Rab-Rab assemblies on two opposing membranes destined to be stably tethered.

To further understand the active “tethering-competent” mode of the Rab G-domains that are closely attached to membrane surfaces, we next examined the effects of lipids on Rab-or Rab G-domain-mediated membrane tethering by performing liposome turbidity assays for full-length Rab5a and HVR-deleted Rab5a (Rab5aΔHVR) with the two different lipid compositions; the physiologically-mimicking complex composition bearing PC, PE, PI, PS, and cholesterol, which was used as the standard in **Figures 2-6** and termed here “complete” (**Figure 7A**), and the non-physiological simple composition bearing PC and PE only, termed “PC/PE” (**Figure 7A**). Strikingly, when tested for full-length Rab5a (final 10 μM, the Rab-to-lipid ratios of 1:100; **Figure 7B**), its intrinsic tethering activity was significantly diminished by omitting two anionic lipid species (PI and PS) and cholesterol from liposomes (**Figure 7B**, left panel), giving the maximum tethering capacities (ΔOD400) of 0.67 ± 0.037 with the complete liposomes but 0.35 ± 0.034 with the PC/PE liposomes (**Figure 7B**, right panel). However, intriguingly, HVR-deleted Rab5a mutant (final 1 μM, the Rab-to-lipid ratios of 1:1,000, **Figure 7C**; final 0.5 μM, the Rab-to-lipid ratios of 1:2,000, **Figure S2**) exhibited almost or completely identical tethering kinetics (**Figure 7C**, left panel; **Figure S2**, left panel) and tethering capacities (**Figure 7C**, right panel; **Figure S2**, right panel) with these two “complete” and “PC/PE” types of liposomes, establishing that the tethering potency of the hyperactive HVR-deleted Rab5a is fully independent of the headgroup composition of lipid bilayers. These results indicate that the HVR linkers, not the G-domains, act as a primary region to interact with the hydrophilic lipid headgroups, thereby guiding the G-domain towards its active tethering-competent state on the membrane surface in the case of Rab5a (**Figure 7B**) or, by contrast, negatively regulating the membrane attachment of the G-domain in the case of Rab4a, which was a quite inefficient membrane tether in its full-length form but found to be a very highly potent tether in the HVR-deleted mutant form (**Figures 2-3**). Furthermore, considering that the HVR deletion allows the Rab G-domain to be in close contact with membrane lipids but simultaneously insensitive to the lipid headgroup composition in the tethering assays (**Figure 7C**), specific hydrophobic interactions between the acyl chains of membrane lipids and the nonpolar surface areas of the Rab G-domains likely induce the proper membrane orientations and suitable structures for achieving rapid and efficient Rab-driven tethering of lipid bilayers.

**Figure 7.**
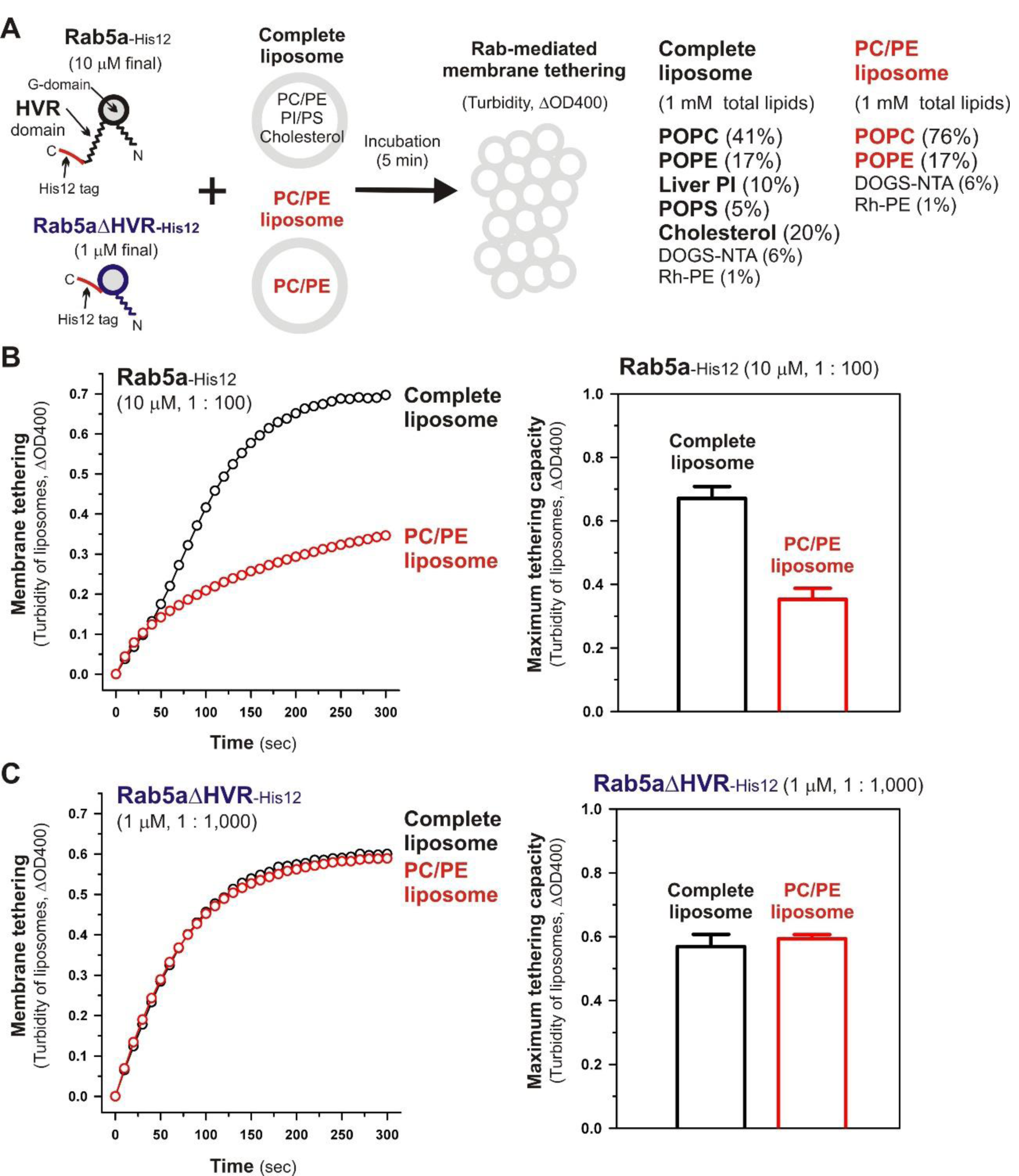
Physiological complex lipid composition is crucial for efficient membrane tethering mediated by full-length Rab5a but not for HVR-deleted Rab5a-mediated membrane tethering. (**A**) Schematic representation of liposome turbidity assays in (**B-C**), testing the requirement of physiological complex lipid composition for Rab5a-mediated membrane tethering. The complete liposomes, which were also used in Figures 2-6, contained PC, PE, PI, PS, and cholesterol, whereas the PC/PE liposomes bore only PC (76%, mol/mol) and PE (17%), in addition to DOGS-NTA and Rh-PE. (**B**) Full-length Rab5a-His12 proteins (final 10 μM) were mixed with the complete liposomes or PC/PE liposomes (200-nm diameter; final 1 mM total lipids) and immediately assayed for the turbidity changes by measuring the optical density at 400 nm (ΔOD400) for 5min (left panel), as in Figure 2. The maximum tethering capacities of full-length Rab5a (right panel) were determined from the kinetic turbidity data, as in Figure 2. The maximum tethering capacities of full-length Rab5a for the complete liposomes are significantly different from those for the PC/PE liposomes (*p* < 0.001, calculated using two-way ANOVA). (**C**) Rab5aΔHVR-His12 proteins (final 1 μM) were mixed with the complete liposomes or PC/PE liposomes and then assayed for the turbidity changes (left panel), as in (**B**) for full-length Rab5a. The maximum tethering capacities of HVR-deleted Rab5a (right panel) were determined from the kinetic data, as in (**B**). Error bars, SD.

### Critical residues in the HVR linker domain of Rab4a for regulating the intrinsic tethering activity

Although full-length, wild-type Rab4a having the HRV linker domain at the C-terminus has been shown to be a very inefficient membrane tether compared to other tethering-competent Rab isoforms such as Rab3a, −5a, −6a, and −7a in a reconstituted system (Inoshita and Mima, 2017; Segawa et al., 2019; **Figure 2, Figure 3**), the present reconstituted tethering assays revealed that deletion of the C-terminal HVR linker allows less-active or inactive Rab4a to be a highly potent membrane tether, exhibiting much higher tethering activity than full-length Rab5a, which was comparable to that of HVR-deleted mutant Rab5a (**Figure 2, Figure 3**). This leads us to postulate that the HVR linker of Rab4a negatively regulates membrane tethering potency of the G-domain via specific amino acid residues within the flexible linker or through simply acting as a wider spacer between the G-domain and the membrane surface, because the Rab4a HVR domain (43 residues long) is longer than the HVRs of the tethering-competent Rab isoforms described above (33-36 residues long) (**Figure 1, Figure 8**; Segawa et al., 2019). To explore this issue, in addition to the full-length (residues 1-218) and HVR-deleted (residues 1-175) forms of Rab4a, we prepared purified proteins of three other Rab4a mutants having the C-terminally-truncated HVR linkers, including Rab4aΔC199 (residues 1-199), Rab4aΔC181 (residues 1-181), and Rab4aΔC177 (residues 1-177) (**Figure 8A-B**), and investigated the tethering activities of these C-terminally-truncated mutants in comparison to full-length and HVR-deleted Rab4a proteins by employing liposome turbidity assays (**Figure 8C-E**).

**Figure 8.**
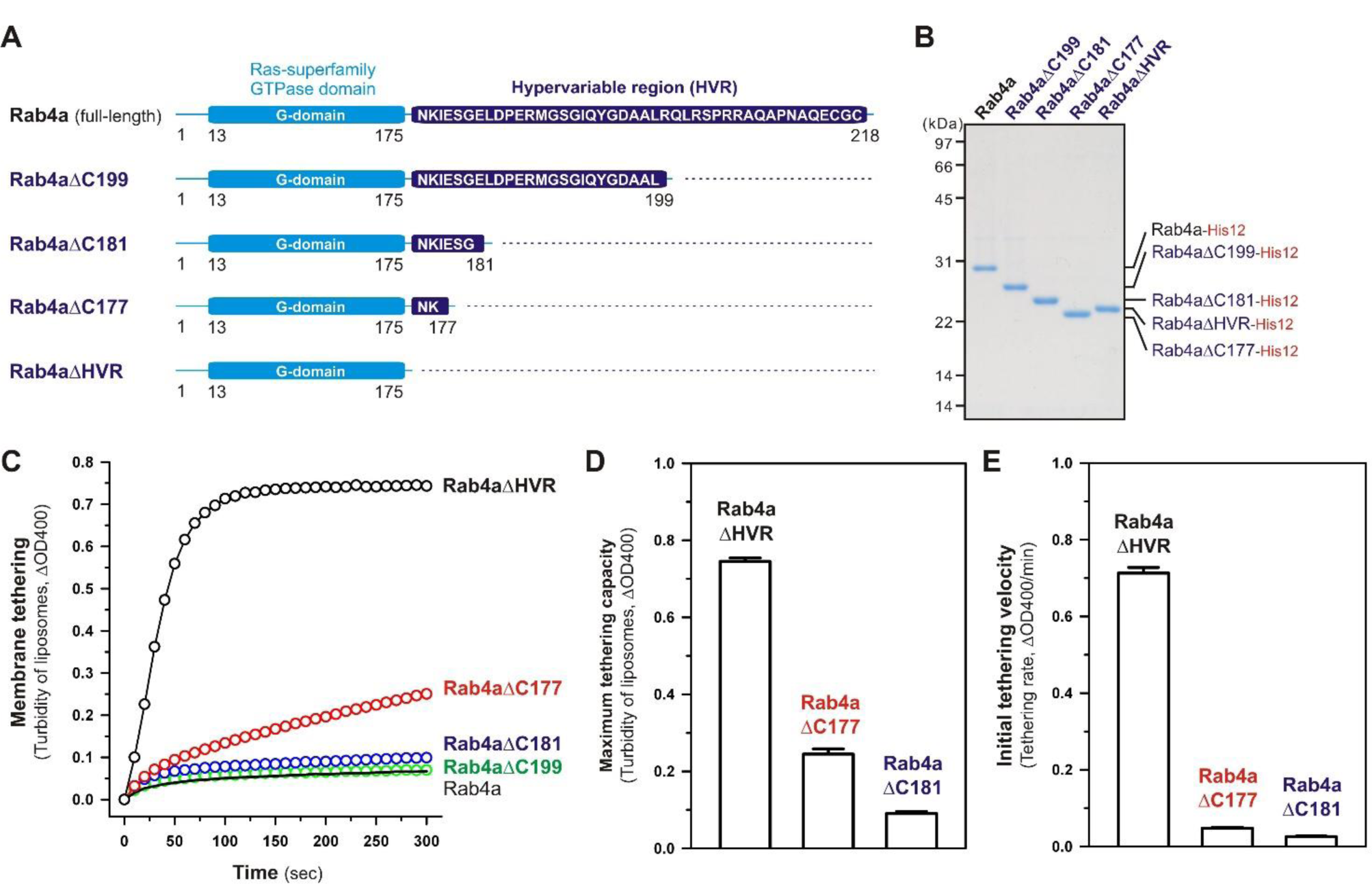
The N-terminal residues in the HVR linker domain are critical to controlling Rab4a-mediated membrane tethering. (**A**) Schematic representation of the C-terminally-truncated Rab4a mutants used for liposome turbidity assays in (**C-E**), including Rab4aΔC199 (Met1-Leu199), Rab4aΔC181 (Met1-Gly181), and Rab4aΔC177 (Met1-Lys177). All of the C-terminally-truncated Rab4a mutant proteins were purified as described for the full-length and HVR-deleted forms of Rab4a. (**B**) The Coomassie blue-stained gel of purified proteins of the C-terminally His12-tagged full-length Rab4a, Rab4aΔC199, Rab4aΔC181, Rab4aΔC177, and Rab4aΔHVR. (**C-E**) Kinetic liposome turbidity assays with the C-terminally-truncated Rab4a mutants. Purified full-length and mutant Rab4a-His12 proteins (final 2 μM) were mixed with DOGS-NTA-bearing liposomes (200-nm diameter; final 1 mM lipids) and immediately assayed for the turbidity changes by measuring ΔOD400 for 5min (**C**), as in Figure 2. The kinetic turbidity data were analyzed by curve fitting to calculate the maximum tethering capacities (**D**) and the initial tethering velocities (**E**), as in Figure 2. The maximum capacities (**D**) and initial velocities (**E**) of Rab4aΔHVR are significantly different from those of Rab4aΔC177 or Rab4aΔC181 (*p* < 0.001, calculated using two-way ANOVA). Error bars, SD.

Among those three Rab4aΔC mutants, Rab4aΔC199 and Rab4aΔC181 remained to be inactive or inefficient in the tethering assays, exhibiting very low tethering capacities and tethering rates similar to those of full-length Rab4a (**Figure 8C-E**), even though the lengths of their truncated HVR linkers (6 or 24 residues long) are significantly shorter than the entire HVR lengths of the tethering-competent Rab isoforms (33-36 residues long). RabΔC177, which has only two extra amino acid residues (Asn-Lys) as a linker between the G-domain and a His12-tag membrane anchor at the extreme C-terminus (**Figure 8A**), appeared to be slightly activated by further truncation of the HVR linker region (**Figure 8C**), but its maximum tethering capacity (0.24 ± 0.014, ΔOD400; **Figure 8D**) and initial tethering velocity (0.048 ± 0.0022, ΔOD400/min; **Figure 8E**) were still substantially lower than those values of the linker-deficient Rab4aΔHVR mutant protein (0.75 ± 0.0089 for the maximum capacity and 0.71 ± 0.015 min^-1^ for the initial velocity; **Figure 8D-E**). These results not only support the idea that the close membrane attachment of the Rab G-domain is required for Rab-driven membrane tethering reactions, but also they suggest that specific amino acid residues in the Rab4a HVR linker domain, particularly several N-terminal residues of the linker, retain the important functions to restrict the intrinsic tethering potency of the Rab4a G-domain, beyond just acting as a non-specific spacer to prevent membrane association of the G-domain.

To obtain further molecular insights into the roles of the N-terminal residues of the Rab4a HRV linker in regulating the tethering potency of the G-domain, we lastly examined the effects of membrane lipids on the tethering activities of the two Rab4aΔC mutants, Rab4aΔC177 and Rab4aΔC181, by employing liposome turbidity assays with the complete liposomes and the PC/PE liposomes lacking two anionic lipids (PI and PS) and cholesterol (**Figure 9**), as tested for full-length and HVR-deleted Rab5a proteins in **Figure 7**. Both of the HVR-truncated Rab4a mutants exhibited significantly higher tethering activities for the PC/PE liposomes than those for the complete liposomes mimicking physiological lipid compositions (**Figure 9**), yielding 2.1-and 3.4-fold increases in the tethering capacities of Rab4aΔC177 (**Figure 9A**) and Rab4aΔC181 (**Figure 9B**), respectively, whereas omission of anionic lipids and cholesterol had little or no effect on the tethering rates and capacities of Rab4aΔHVR that lacks the entire HRV linker (**Figure S3**). These results suggest that the two (Asn-Lys) or six (Asn-Lys-Ile-Glu-Ser-Gly) N-terminal residues in the Rab4a HRV linker domain contribute to limiting the intrinsic tethering potency of the G-domain through their interactions with the polar lipid headgroups. Considering that recent computational studies by molecular dynamics simulations of membrane-bound K-Ras, belonging to the Ras superfamily, reported the multiple orientation states of the K-Ras G-domain on the membrane surface that are affected by the C-terminal HVR linker and anionic membrane lipids (Prakash et al., 2016; Prakash and Gorfe, 2019; Neale and Garcia, 2020; Ngo et al., 2020), it is conceivable that the hydrophilic interactions between the N-terminal residues of the Rab4a HVR linker and anionic lipids may guide the G-domain into the specific “tethering-incompetent” membrane orientation.

**Figure 9.**
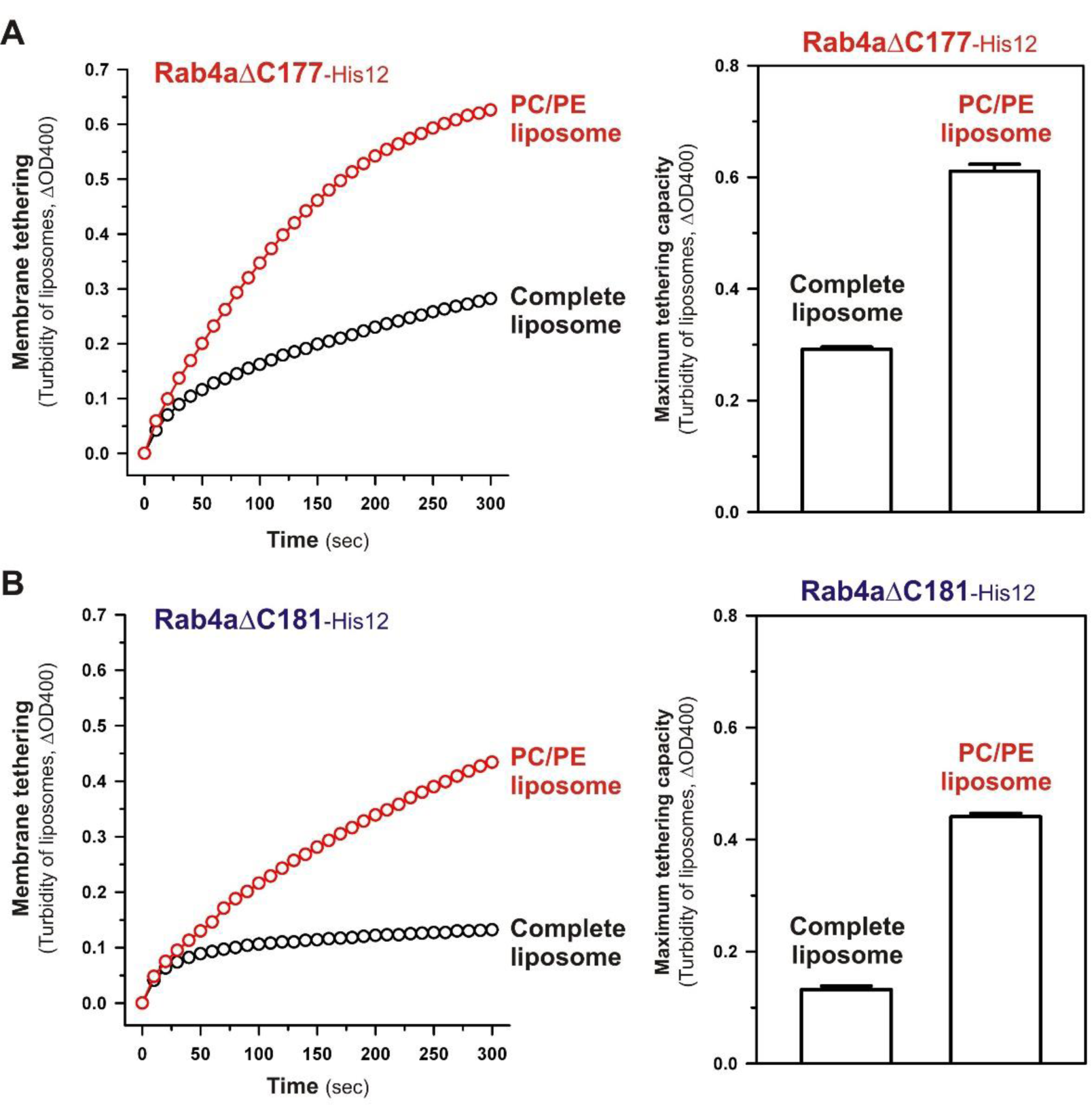
The C-terminally-truncated Rab4a mutants, Rab4aΔC177 and Rab4aΔC181, exhibit higher membrane tethering potency for the non-physiological PC/PE liposomes than that for the physiologically-mimicking complete liposomes. (**A**) Kinetic liposome turbidity assays for Rab4aΔC177 with the complete liposomes containing PC, PE, PI, PS, and cholesterol and the PC/PE liposomes containing PC and PE only. Purified Rab4aΔC177-His12 proteins (final 2 μM) were mixed with the complete liposomes or PC/PE liposomes (200-nm diameter; final 1 mM lipids) and immediately assayed for the turbidity changes by measuring ΔOD400 for 5 min (left panel), as in Figure 7. The maximum tethering capacities of Rab4aΔC177 (right panel) were determined from the kinetic turbidity data, as in Figure 7. The maximum tethering capacities of Rab4aΔC177 for the complete liposomes are significantly different from those for the PC/PE liposomes (*p* < 0.001, calculated using two-way ANOVA). (**B**) Kinetic liposome turbidity assays for Rab4aΔC181 with the complete liposomes and PC/PE liposomes. Purified Rab4aΔC181-His12 proteins were mixed with the complete liposomes or PC/PE liposomes and then assayed for the turbidity changes (left panel), as in (**A**) for Rab4aΔC177. The maximum tethering capacities of Rab4aΔC181 (right panel) were determined from the kinetic turbidity data, as in (**A**). The maximum tethering capacities of Rab4aΔC181 for the complete liposomes are significantly different from those for the PC/PE liposomes (*p* < 0.001, calculated using two-way ANOVA). Error bars, SD.

### Conclusions

By comprehensively and quantitatively investigating the intrinsic membrane tethering potency of human endosomal Rab-family small GTPases (Rab5a, Rab4a) and their mutant forms lacking the C-terminal HVR domains (Rab5aΔHVR, Rab4aΔHVR) or having the truncated HVR linkers (Rab4aΔC mutants) in a chemically-defined reconstitution system (**Figures 2-9**), the present studies provide novel insights into the mechanistic basis of membrane tethering reactions driven by Rab small GTPases: (1) Close attachment of the globular G-domains to membrane surfaces is a vital step to fully activate the intrinsic potency of Rab proteins to trigger selective *trans*-Rab-Rab assemblies and subsequently drive efficient membrane tethering reactions; (2) the HVR linkers control, either positively or negatively, the close membrane attachment of the G-domains through the interactions between the polar residues in HVRs and the headgroups of membrane lipids; and (3) the active “tethering-competent” state of the G-domains closely attached onto membrane surfaces is completely insensitive to the composition of lipid headgroups, suggesting the importance of the hydrophobic interactions between the G-domain surfaces and the non-polar regions of membrane lipids. Finally, based on the current findings, future studies will focus on deciphering the protein-protein and protein-lipid interfaces in the Rab G-domains during *trans*-Rab-Rab assembly and Rab-mediated membrane tethering.

## Author contributions

JM designed the research. JM, SU, and NT performed the experiments. JM, SU, and NT analyzed the data. JM wrote the manuscript.

## Acknowledgements

We thank Megumi Shinguu and Kazuya Segawa (Institute for Protein Research, Osaka University) for their contributions to preparation of recombinant proteins of human Rabs. We are grateful to Dr. Genji Kurisu and Dr. Hideaki Tanaka (Institute for Protein Research, Osaka University) for access to dynamic light scattering measurements. This study was in part supported by the Program to Disseminate Tenure Tracking System from the Ministry of Education, Culture, Sports, Science and Technology, Japan (MEXT) and Grants-in-Aid for Scientific Research from MEXT (to JM).

## Supplemental Figures

**Figure S1.**
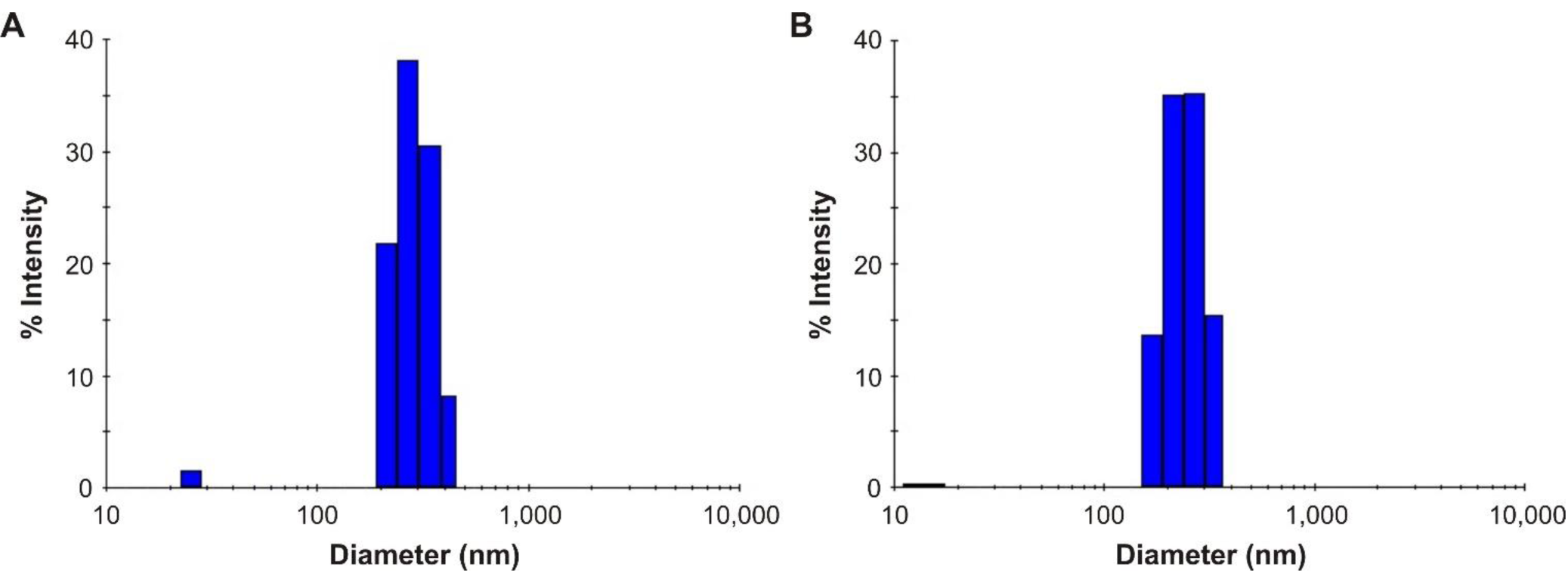
Dynamic light scattering measurements of synthetic liposomes used in the present reconstitution studies. (**A-B**) Histograms of size distributions of typical liposome preparations, the Rh-labeled 200-nm complete liposomes containing PC, PE, PI, PS, and cholesterol (**A**) and the Rh-labeled 200-nm PC/PE liposomes containing PC and PE (**B**), measured by dynamic light scattering.

**Figure S2.**
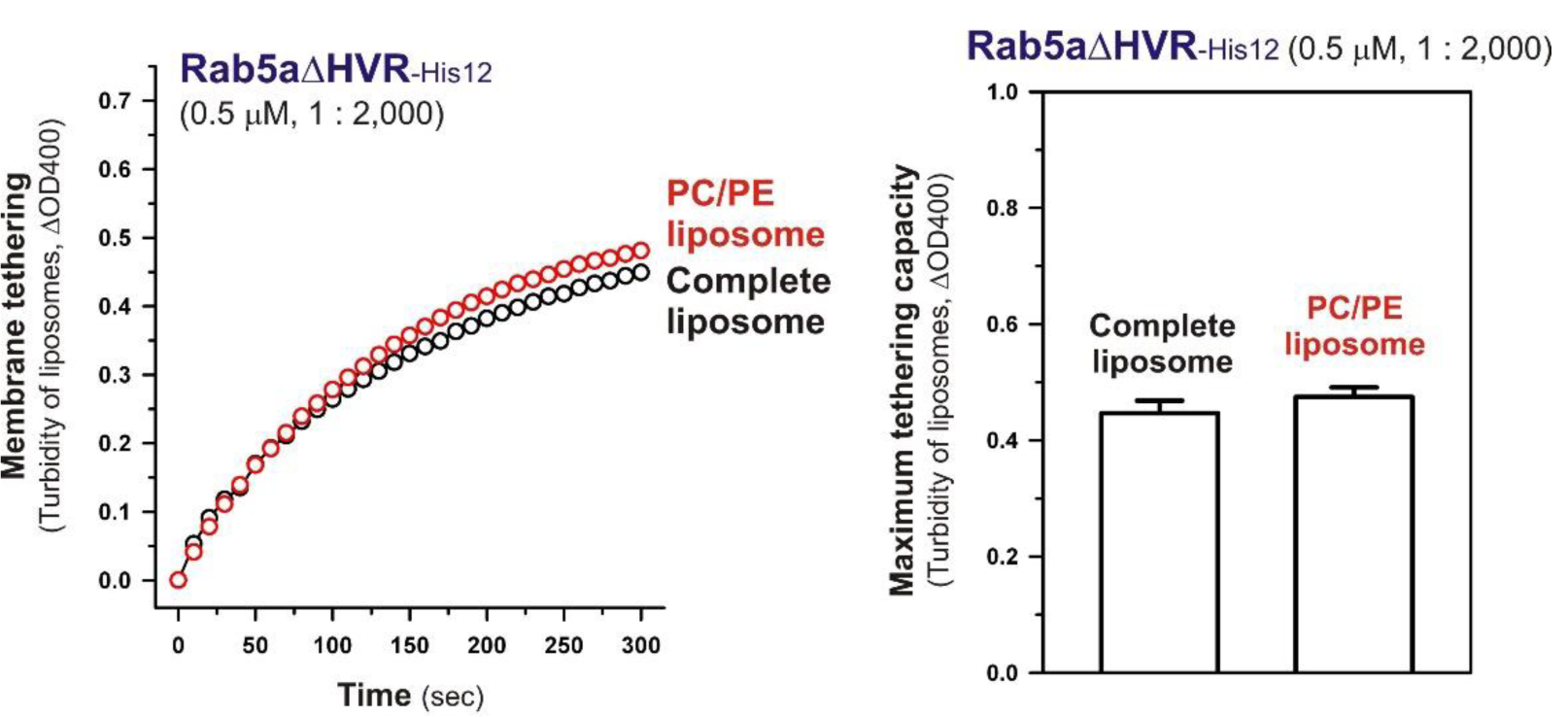
Liposome turbidity assays for Rab5aΔHVR-mediated membrane tethering with the complete liposomes and PC/PE liposomes, testing at the lower Rab-to-lipid molar ratios of 1:2,000. Rab5aΔHVR-His12 proteins (final 0.5 μM) were mixed with the complete liposomes or PC/PE liposomes (200-nm diameter; final 1 mM lipids) and then assayed for the turbidity changes (left panel) as in Figure 7C, but at the Rab-to-lipid molar ratios of 1:2,000. The maximum tethering capacities of Rab5aΔHVR (right panel) were determined from the kinetic data, as in Figure 7C. Error bars, SD.

**Figure S3.**
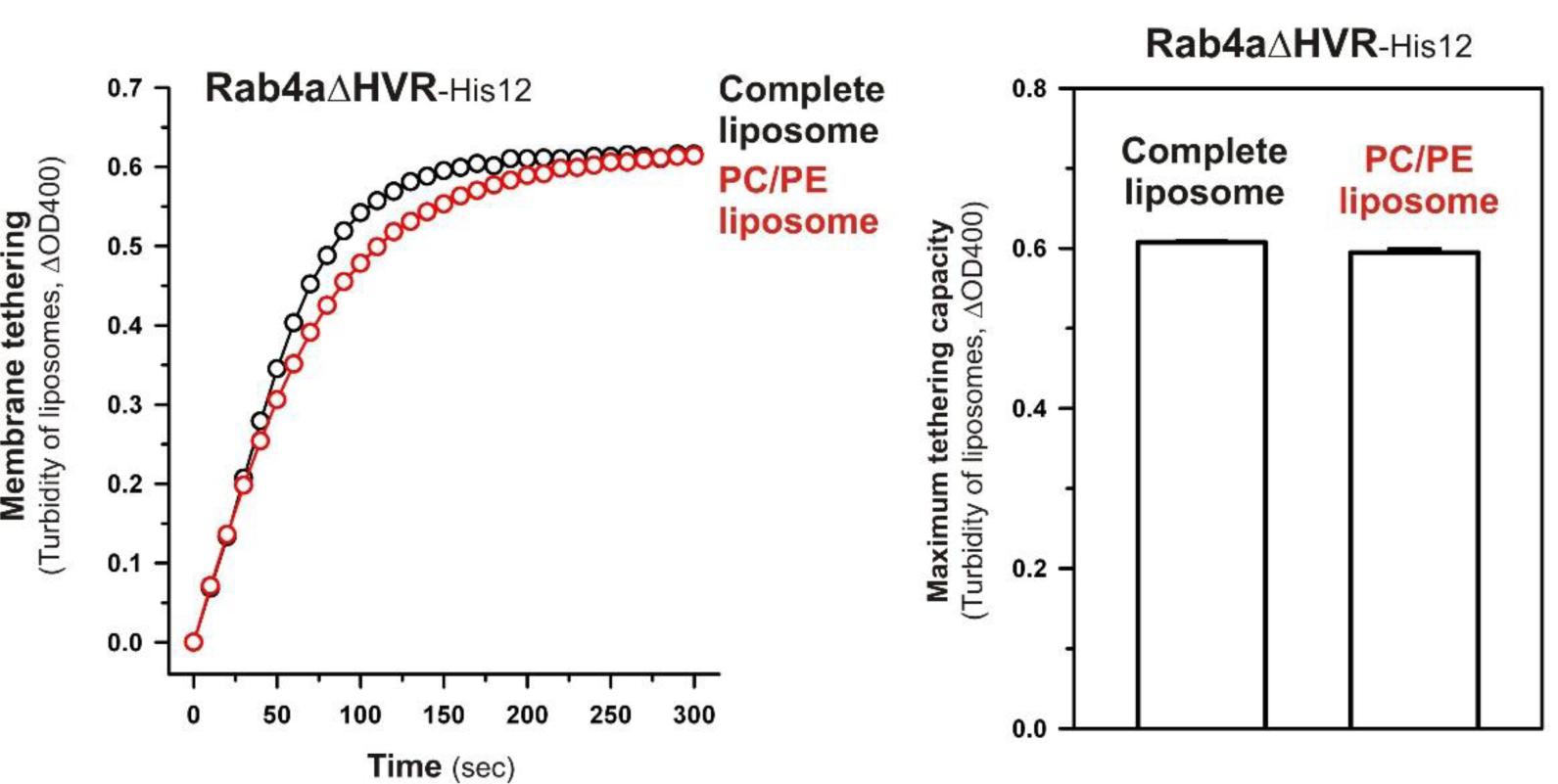
Liposome turbidity assays for Rab4aΔHVR-mediated membrane tethering with the complete liposomes and PC/PE liposomes. Rab4aΔHVR-His12 proteins (final 1 μM) were mixed with the complete liposomes or PC/PE liposomes (200-nm diameter; final 1 mM lipids) and then assayed for the turbidity changes (left panel), as in Figure 7 and Figure 9. The maximum tethering capacities of Rab4aΔHVR (right panel) were determined from the kinetic data, as in Figure 7 and Figure 9. Error bars, SD.

